# TRACKING CLONAL AND PLASMID TRANSMISSION IN COLISTIN AND CARBAPENEM RESISTANT *KLEBSIELLA PNEUMONIAE*

**DOI:** 10.1101/2024.08.20.608806

**Authors:** Ifeoluwa Akintayo, Marko Siroglavic, Daria Frolova, Mabel Budia Silva, Hajo Grundmann, Zamin Iqbal, Ana Budimir, Sandra Reuter

## Abstract

The surveillance of mobile genetic elements facilitating the spread of antimicrobial resistance genes has been challenging. Here, we tracked both clonal and plasmid transmission in colistin- and carbapenem-resistant *K. pneumoniae* using short and long read sequencing technologies. We observed three clonal transmissions, all containing IncL plasmids and *bla*_*NDM*−1_, although not co-located on the same plasmid. One IncL-*bla*_*NDM*−1_ plasmid had been transferred between ST392 and ST15, and the promiscuous IncL-*bla*_*OXA*−48_ plasmid was likely shared between a singleton and a clonal transmission of ST392. Plasmids within clonal outbreaks and between clusters and STs had 0-2 SNP differences, showing high stability upon transfer to same or different STs. The simplest explanation of a single common IncL-*bla*_*NDM*−1_ plasmid spreading was in fact false, and we found *bla*_*NDM*−1_ in the context of five different plasmids, emphasizing the need to investigate plasmid-mediated transmission for effective containment of outbreaks.

**IMPORTANCE:** Antimicrobial resistance occupies a central stage in global public health emergencies. Recently, efforts to track the genetic elements that facilitate the spread of resistance genes to determine so-called plasmid outbreaks have been described, however, such short read sequencing hinders full knowledge about plasmid structure and makes this approach very challenging to implement. In this study we used both short and long read sequencing in clinical Klebsiella pneumoniae from University Hospital Centre Zagreb, Croatia which were resistant to both last resort antibiotics colistin and carbapenem. Our results show complex transmission networks and sharing of plasmids, emphasizing multiple transmissions of plasmids harbouring carbapenem and/or colistin resistance genes between and within K. pneumoniae clones. Only full length sequencing plus a novel way of determining plasmid clusters resulted in the full picture, showing how future active monitoring of plasmids as a vital tool for infection prevention and control could be implemented.

## INTRODUCTION

Antimicrobial resistance is one of the biggest challenges in modern medicine and a major concern to public health worldwide (1). The European Centre for Disease Prevention and Control (ECDC) reports high and increasing antibiotic resistance among Gram-negative bacteria in many parts of Europe, including Croatia (2),one of which is *Klebsiella pneumoniae* – a major cause of hospital-acquired infections. In recent years, carbapenem-resistant *K. pneumoniae* (CRKP) lineages have become pathogens of major concern, especially in the healthcare system (3) owing to their ability to acquire resistance to last resort antimicrobials, like carbapenems and colistin, thereby making infections particularly difficult to treat (4). Resistance to carbapenems can be mediated by mutations in genes that lead to overexpression of efflux pumps or to porin deficiency, along with expression of AmpC beta-lactamases or extended spectrum beta-lactamases (ESBL). However most worrying is resistance due to acquisition of genes encoding carbapenem-hydrolyzing enzymes (i.e. carbapenemases) (5). The major carbepenemases implicated in resistance to common carbapenems among Enterobacterales are *bla*_*KPC*_, *bla*_*OXA*−48_, *bla*_*NDM*−1_ and *bla*_*VIM*_ (6). Therapeutics deployed for the treatment of multidrug and CRKP infections involve the use of antibiotics of last resort, such as colistin (polymyxin E) (7). As with carbapenem resistance, colistin resistance (ColR) is on the rise worldwide, and has been reported in several parts of Europe, including Croatia (8). Mechanisms of bacterial resistance to colistin involve lipopolysaccharide modifications, mediated by mutations on chromosomal genes (mgrB, phoPQ and pmrAB), and acquisition of mobile colistin resistance (mcr) genes, which have been reported in *K. pneumoniae* (9). Until now, different variants of the mobile colistin resistance gene mcr (*mcr*-1 to *mcr*-10) have been identified in several bacterial species from various sources (10). The rapid dissemination of carbapenem resistance genes is most often driven by mobile genetic elements such as plasmids, which are transferable within and between bacteria species or strains (11). Outbreaks that are plasmid-associated are increasingly studied and have been reported to facilitate the spread of antimicrobial resistance genes in multiple bacterial strains or species, especially in clinical settings (12-13). Hence, outbreak investigations of carbapenem and colistin resistant strains and their mobile elements are critical to understand epidemiology and efficiently strategize infection prevention and control measures. In this study, we investigated an outbreak of colistin and carbapenem-resistant *K. pneumoniae* (ColR-CRKP) strains in a university hospital in Zagreb, Croatia. Short and long read sequences were used to investigate and differentiate clonal and plasmid mediated transmission of antimicrobial resistance.

## MATERIALS AND METHODS

### Culture, DNA extraction and whole-genome sequencing

A total of 46 *K. pneumoniae* isolates from the University Hospital Centre (UHC) Zagreb, Zagreb, Croatia, were identified as part of routine surveillance and clinical testing of microbiological samples from all clinical departments. Clinical samples comprise 54% of tested samples. Samples were cultured on Columbia blood agar containing 5% sheep blood. Isolates identification was confirmed by MALDI-TOF mass spectrometry (Bruker Microflex LT, Bremen, Germany). Antimicrobial susceptibility was determined by VITEK 2 compact system (bioMérieux, Paris, France) and interpreted according to the European Committee on Antimicrobial Susceptibility Testing (EUCAST) criteria (available at http://www.eucast.org/clinical_breakpoints/). Information on all samples can be found in supplementary table S1. These samples were then transferred to the Medical Center – University of Freiburg, Freiburg, Germany for whole genome sequencing (WGS). Species identification was confirmed using MALDI-TOF mass spectrometry (BRUKER), and isolates were cultured overnight at 37°C on blood agar. DNA was extracted according to manufacturer’s instructions using a high pure PCR template preparation kit (Roche diagnostics). All isolates were short-read sequenced with an Illumina MiSeq using Nextera DNA flex library preparation V2 300 cycle PE kit according to manufacturer’s instructions and all isolates were long-read sequenced with Oxford Nanopore Technology (ONT) GridION platform using the ligation protocol SQK-LSK109 with native barcode EXP-NBD104 kit and SQK-LSK 114 (Oxford Nanopore Technology, Oxford, UK) in accordance with the manufacturer’s protocol. The sequencing was performed using FLO-MIN106 (v.R9.4) and FLO-MIN114 (v.R10.4.1) flow cell type.

### Quality control of sequence analysis

De novo assemblies of the Illumina sequence reads were generated using SPAdes v3.13.1 (http://cab.spbu.ru/software/spades/) with kmer sizes 21, 33, 55, 77, 99, 109, and 123 (14). Assemblies were then filtered to only include contigs with a minimum of 500bp. The sequence type was determined using multi-locus sequence typing (MLST) v2.10 (15). All sequence reads were mapped to the reference genome MGH78578/ATCC700721 (CP000647) using SMALT v0.7.6 (16) and a taxonomic check was carried out using kraken (17). Quality control (QC) criteria include read allocation to the respective expected genus, coverage greater than 30X, expected genome size, number of contigs less than 500, largest contig greater than 100,000bp, N50 greater than 100,000bp, and identification of MLST alleles (Supplementary Table S1).

ONT fast5 generated output file was basecalled on the GridION using guppy v5.0.12. Sequence reads were trimmed using porechop v0.2.4 (18) to remove adapters from read ends and within the reads, using default parameters. The trimmed reads were filtered to retain long reads using filtlong v0.2.1 (19). Hybrid assemblies were generated using unicycler v0.5.0 (20), and contigs with less than 500bp were excluded. QUAST v5.0.2 (21) was used to generate the assembly statistics.

All sequencing statistics, as well as accession numbers for raw reads and assemblies can be found in supplementary table S1. Sequencing data was deposited under ENA project PRJEB76496.

### Phylogenetic analysis and isolate characterization

Single nucleotide polymorphisms (SNPs) were filtered from the mapping data with GATK (22), the variant filtered files were converted to a fasta file, where SNP sites and absent sites (N) were replaced in the reference genome. An alignment file was generated, and mobile genetic elements were removed (https://github.com/sanger-pathogens/remove_blocks_from_aln). Gubbins was employed to detect recombination events and reconstruct a phylogenetic tree (23). Trees were visualized using figtree v1.4.4 (https://github.com/rambaut/figtree) and iTOL v6.5 (24). Antimicrobial resistance genes and point mutations were identified from the assembled files using AMRFinderplus and its NCBI database (25).

### Plasmid analysis and visualization

Plasmid replicons were predicted using abricate and PlasmidFinder database (26). The plasmid contigs were extracted from the hybrid assembled files and annotated using bakta (27). Plasmids were clustered in two independent ways. First, they were clustered using MASH v1.4.5 (28), using a Mash distance of threshold of 0.001 to define highly similar plasmid sequences as previously described (29). However this approach is not a good estimator of structural relatedness (e.g “these plasmids differ by one deletion and one inversion”), so for this we clustered the plasmids using pling v1.0.1 with default parameters. The output is a relatedness network where nodes are plasmids, and edges are labelled with two numbers: a “containment distance” (what proportion of the smaller plasmid is not alignable to the larger) and a rearrangement distance (how many rearrangements/indels separate these two plasmids) (50). Similarities searches were performed using NCBI BLAST (30), linear comparative analyses of the plasmid were generated using Easyfig v2.2.2 (31) and circular plot of the novel plasmid was generated using DNAplotter (32). IncL plasmids were downloaded from the plasmid database (PLSDB) (33) and together with the IncL plasmids in this study, the sequences were compared pairwise using two similarity metrics: (i) Mash similarity and (ii) gene content Jaccard similarity as described in a previous study (34) and the figure was generated using cytoscape (35).

## RESULTS

### Basic overview of the population

We included 46 isolates ColR-CRKP from the University Hospital Centre (UHC) Zagreb, isolates over a nearly two-year period. We identified two major sequence types, ST15 (n=29) and ST392 (n=8), with other isolates belonging to ST101 (n=4), ST17 (n=1), ST147 (n=1), ST273 (n=1), ST274 (n=1) and ST5081 (n=1) (Fig. 1a). Genetic information based on SNP distance did not indicate a clonal transmission cluster in ST101. The metadata showed that the patients were not at any point in time in the same ward, hence we focused our clonal transmission analysis on ST392 and ST15.

**Figure 1:**
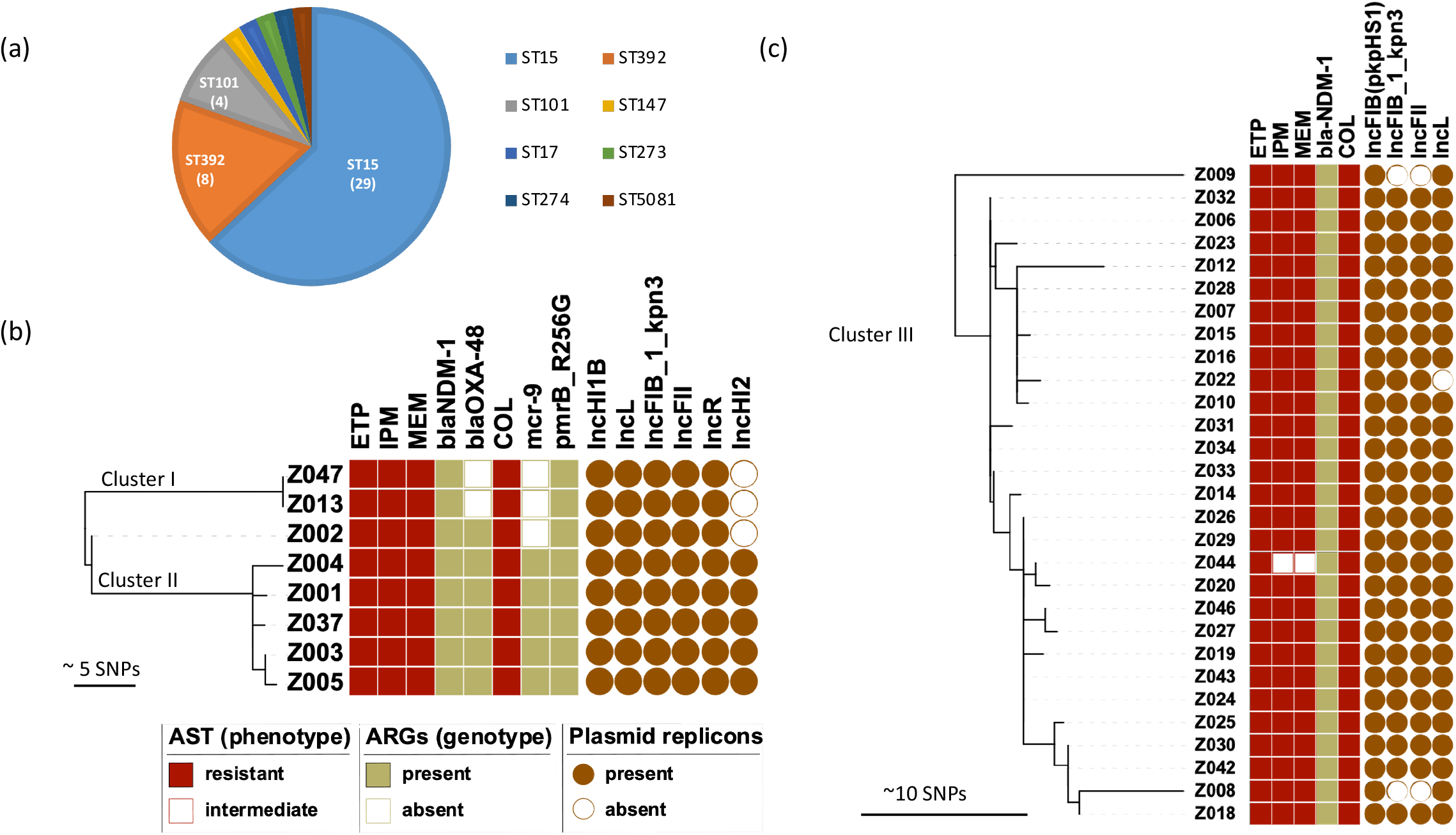
Overview of isolates and phylogenetic reconstruction of main STs. (a) Distribution of the sequence types (STs). Phylogeny of (b) ST392 and (c) ST15. Results of phenotypic testing for carbapenem and colistin resistance are followed by their corresponding genotype. Distribution of main detected plasmid incompatibility groups are shown. ETP, ertapenem; IPM, imipenem; MEM, Meropenem; COL, colistin.

### Clonal transmission patterns of ST392 and ST15

We determined genetic relatedness of ST392 isolates (Fig. 1b). The phylogeny showed two distinct clusters (Fig. 1b) that present putative separate transmission chains separated by 14-22 SNPs. There was a singleton isolate Z002 distinct from all the others by 7-8 SNPs. Cluster I involved 2 patients (Z047 and Z013) with 0 SNPs apart while cluster II involved 5 patients with 1-4 SNPs apart. Spatial and temporal relationship of the patients involved in each cluster showed that they were at some point in the same ward at the same time (Fig. 2). Cluster II patients overlapped in an internal medicine ICU ward between September and December 2021 while cluster I patients overlapped in a surgical ICU ward in February 2022. Isolate Z004 was temporally the first isolate detected, before other patients entered the internal medicine ICU, and can therefore be considered the index patient of cluster II. Isolates in this cluster were highly non-susceptible and showed phenotypic pandrug resistance to nine classes of antibiotics, including polymyxins and carbapenems (Supplementary Table S1). The other isolates were non-susceptible except to fosfomycin (Supplementary Table S1). A mutation in the pmrB gene (pmrB_R256G) was the only colistin resistance determinant detected in cluster I while cluster II had both pmrB_R256G and *mcr*-9. Carbapenemase *bla*_*NDM*−1_ was detected in all ST392 isolates, with cluster II and Z002 additionally carrying *bla*_*OXA*−48_. A full set of detected resistance genes can be found in Supplementary Table S1. A total of 5 plasmid replicons were detected, including IncHI1B, IncFII, IncFIB, IncR, and IncL with an additional plasmid replicon IncHI2 only detected in cluster I (Fig 1b).

**Figure 2:**
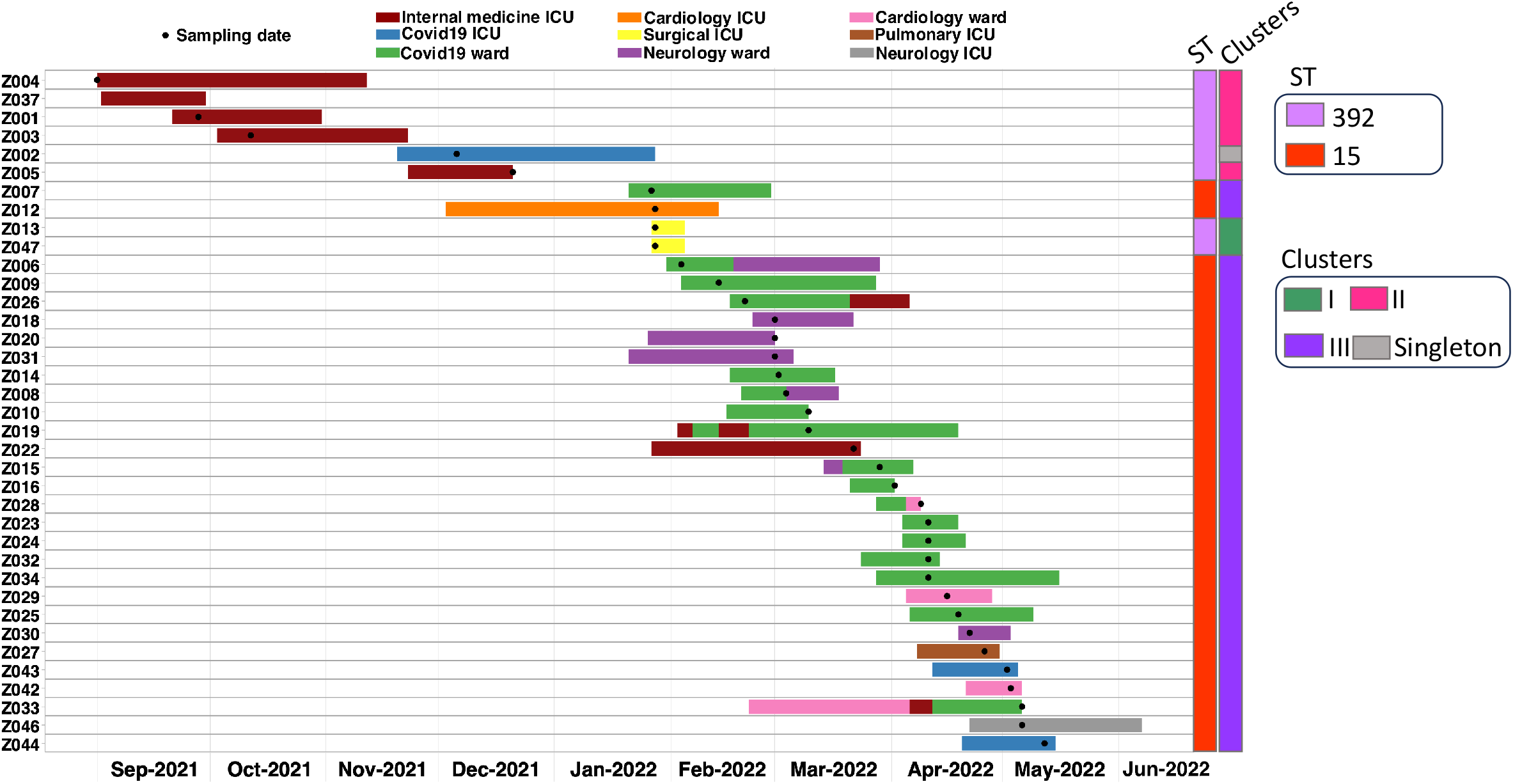
Spatial and temporal relationship of the patients within the hospital for the two main STs and clonal transmission clusters. Patients are sorted by the isolation date of their ColR-CRKP, with the stay on a ward depicted by different colours. Identified ST and cluster allocation is given.

Looking at ST15 (Fig. 1c, Fig. 2), we identified 29 patients in 7 different wards over 6 months, thus the phylogeny showed more genetic diversity. Except for isolate Z009 which is 15-22 SNPs apart, the other isolates had 0-11 SNPs difference, mean and median 3 SNPs. There were two main wards involved in this outbreak, a COVID-19 ward and a neurology ward. A number of patients only had temporal overlaps but no direct ward contact with other patients of this cluster (Z012, Z030, Z027, Z043, Z046, Z044; Fig. 2), however their position within the genetic diversity within this outbreak would suggest they are part of it (Fig. 1c). As with ST392, ST15 was also highly non-susceptible with phenotypic resistance towards six classes of antibiotics (Fig. 1c, Supplementary Table S1). The only carbapenemase detected was *bla*_*NDM*−1_. No known colistin resistance determinant was detected amongst the isolates (Supplementary Table S1). Four different plasmid replicons were found, mainly IncFIB(pkpHS1), IncFIB_1_kpn3, IncFII, and IncL (Fig. 1c).

### Plasmid makeup and transmission of IncL-type plasmids

The variation in the genotypic resistance pattern and plasmid types in this collection prompted further investigation into the carbapenemases and the plasmids carrying them. We detected common replicon types and common resistance genes, namely IncL plasmids and *bla*_*NDM*−1;_ however, the exact makeup of these plasmid was unknown, as was the location of *mcr*-9.

The IncL plasmid replicon was seen in ST15 excluding Z022, all isolates of ST392, and ST5081, but from comparison analysis based on mash distances (Figure 3a), clade 1 comprises IncL plasmids found in ST392-cluster II and ST392 singleton Z002 and clade 2 comprises IncL plasmids found in isolates of ST392-cluster I, ST15 (cluster III), and ST5081. The plasmids in clade 1 had a size of 64 kb and carried only the *bla*_*OXA*−48_ gene, hence we will refer to it as IncL-64kb. The plasmid sequence was compared to the public database using BLAST and was found to be the promiscuous, well-documented pOXA-48 plasmid that is carried in *K. pneumoniae, E. coli*, and other Enterobacterales (Fig.3b). Since this plasmid is highly conserved, with little variation in size and few mutations, within the ST392-cluster II and the singleton isolate we cannot distinguish between the uptake of this plasmid from a common source or plasmid exchange between these two. The IncL plasmid in clade 2 had a size of 96 kb and carried the *bla*_*NDM*−1_ gene with other genes that confer resistance to aminoglycosides, chloramphenicol, sulfonamides and quinolones. In this study, we refer to this plasmid as IncL-96kb. When compared to the public database using BLAST, we found only partial query coverage (68% and 64% to accessions OW970501 and OW969621, respectively) corresponding to the typical pOXA-48 IncL plasmid regions but with a deletion of *bla*_*OXA*_−48. These plasmids also formed their own cluster when compared with other IncL plasmids in PLSDB (Fig. 3c). Both IncL-64kb and IncL-96kb plasmids thus shared a common backbone including the conjugational transfer system, with differences found in the AMR regions (Fig. 3d; Supplementary Fig. 2). The novel IncL-96kb plasmid showed 0-2 SNPs within clusters and upon the transfer between STs, with only an insertion of an additional insB insertion element and hypothetical protein in ST15 and 5081, respectively. The occurrence of such a distinct and unique novel IncL-96kb plasmid and high degree of conservation suggests that it is spreading between the ST392-cluster I, ST15-cluster III and ST5081-Singleton within the hospital.

**Figure 3:**
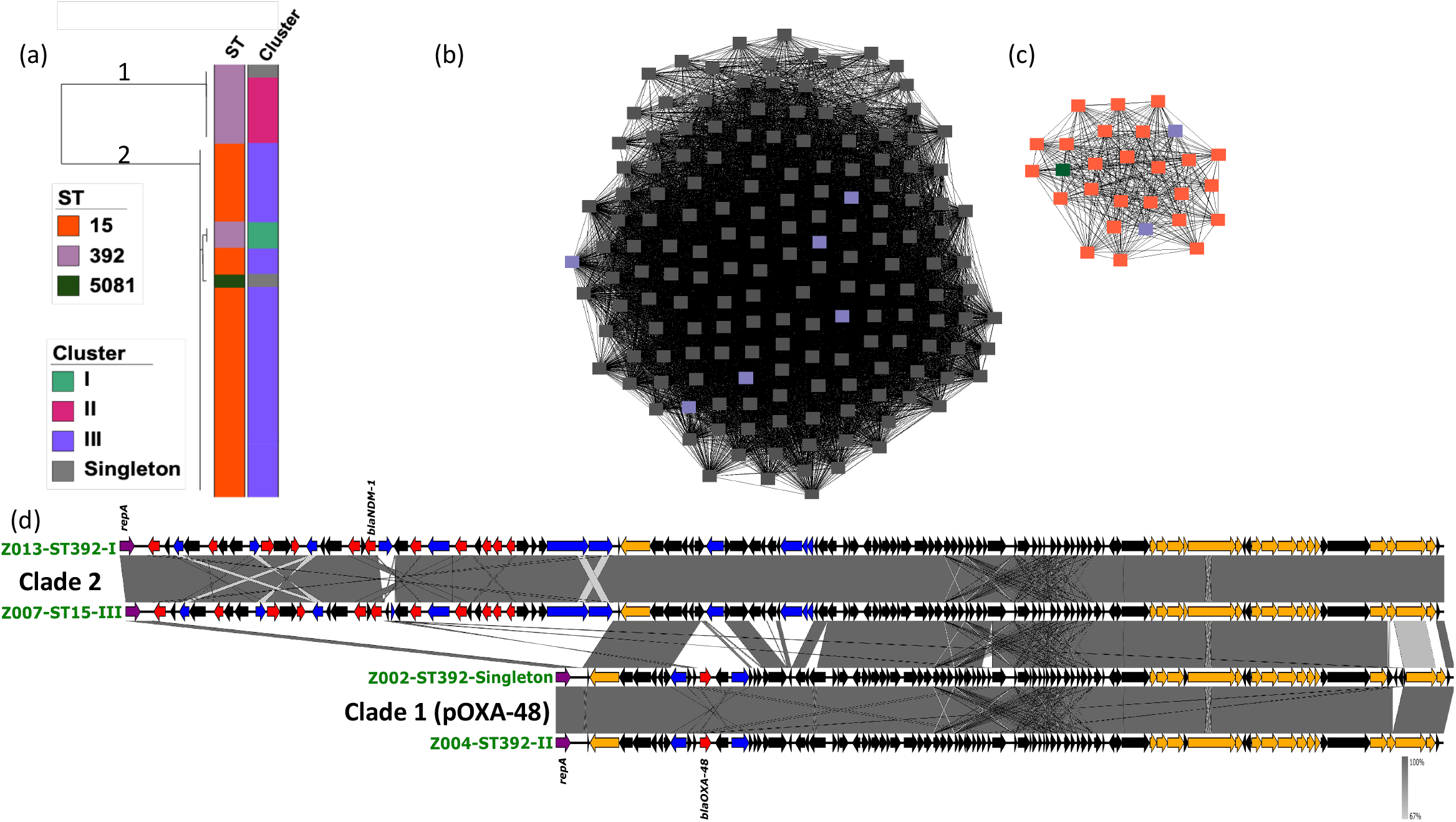
Comparative genomics analysis of IncL plasmid. (a) Mashtree showing IncL plasmid clustering. Clade 1 is IncL-64kb from ST392-cluster II and singleton Z002 while clade 2 is IncL-96kb in ST15 referred to as cluster III and ST392-cluster I. (b) IncL plasmid clustering using mash similarity and gene content jaccard similarity for IncL-64kb and (c) IncL-96kb. (b) Black nodes are IncL pOXA-48 in the plasmid database (PLSDB) while (c) the IncL-96kb do not cluster to any plasmid in the PLSDB. The colours correspond to Sts as in (a). All overlapped nodes were merged. (d) Linear map showing the genetic content of the plasmids from 2 representatives from each clade, Clade 1 (IncL-64kb pOXA-48) and Clade 2 (IncL-96kb). The grey area indicates regions of shared similarities. Red arrows indicate resistance genes, blue arrow indicates transposons, orange arrows indicate the conjugal transfer genes and black arrows are either hypothetical proteins or genes of the replication system.

### Characterization of the mobile genetic context of *bla*_*NDM*_−1 and *mcr*-9

ST392-cluster II isolates contained an IncHI2 *bla*_*NDM*−1_/*mcr*-9 plasmid with sizes ranging from 276 kb to 281 kb (Supplementary Table S2). Other genes conferring resistance to aminoglycosides (*aac-(3)-IIg, aac-(6)-IIc*), beta-lactam (*bla*_*OXA*−1_, *bla*_*TEM*−1_), chloramphenicol (*catB3*), sulphonamides (*sul1*), trimethoprim (*dfrA-19*) and quinolones (*qnrB1*) were also carried. The IncHI2 plasmids were highly similar, having 96-100% coverage and 99% identity with each other. After mapping Illumina short reads of all ST392 cluster II isolates against the hybrid assembly of the longest plasmid, the plasmids were 0-2 SNPs apart with mash distance ranging from 1.4e-05 to 1.4e-03. From the comparative analyses, we observed two small regions differing in isolate Z005 accounting for the observed size difference (Supplementary Fig. 3b-d), the first of which was potentially by IS-mediated excision since it is flanked by repeats. This patient was the last sampled, with the isolate collected two and half months after the previous sample. The plasmid was compared to the public database using BLAST and found to be closely related to a *mcr*-9 harbouring IncHI2 plasmid (accession no. CP030742.1) found in another *K. pneumoniae* isolate, although this plasmid did not contain *bla*_*NDM*_−1.

In contrast, the *mcr*-9 and *bla*_*NDM*−1_ genes detected in singleton ST274 (Z036) were carried on separate plasmids, IncHI2 and IncC, respectively. Comparative structural analysis of the IncHI2 plasmids using pling revealed that previously described five plasmids in ST392 cluster II are almost identical structurally, however the other IncHI2 plasmid (from Z036) is considerably different, separated by at least 12 structural events (Supplementary Fig. 3a). The region carrying the *bla*_*NDM*−1_ gene is absent. Aside from that, the Z036 IncHI2 plasmid had a large inversion with further genes lost (Supplementary Fig 3b). The relatedness of these IncHI2 plasmids and absence of similar plasmids previously described could mean that this plasmid has also been shared between ST392-cluster II and Z036 (ST274) with subsequent genetic modifications, however, given the higher number of genetic changes required and the fact that Z036 was isolated before ST392-cluster II, this seems less likely.

The IncC plasmid (175kb) carrying the *bla*_*NDM*−1_ gene in Z036 had similarities (Mash distance 1.4e-03) with the IncC plasmid (163 kb) carrying *bla*_*NDM*−1_ found in Z011 (ST101 singleton) except for a region lost in Z011 containing qnrA6 bounded by IS element and hypothetical proteins (Supplementary Fig. 4). When compared to the public database, we found identical IncC *bla*_*NDM*−1_ plasmids in *E. coli* and *K. pneumoniae* (100% query coverage and percentage identity; MG450360.1, CP030744.1). This again makes it impossible to distinguish between a possible plasmid transfer between these STs and independent uptake from a shared source.

### Multiple plasmids carrying carbapenemases in other STs

The remaining isolates all had one or two carbapenem resistance gene(s) carried on a variety of plasmids (Supplementary Table S2). *bla*_*OXA*−48_ was seen in all ST101 isolates (Z011, Z021, Z038 and Z039) and was carried on an IncR-FIA plasmid. Apart from Z038, the other IncR-FIA plasmids clustered together, having 100% shared sequence content, but are separated by 5-7 rearrangement events (Supplementary Fig. 5). When all the IncR-FIA plasmids were compared to the public database using BLAST, we found 99-100% query coverage and percentage identity to IncR-FIA plasmids (CP083022.1 and MN218814.1) carrying *bla*_*OXA*−48_ found in *K. pneumoniae* isolated in Switzerland and Serbia, respectively, in the year 2017.

Z045(ST147) carried a *bla*_*NDM*−1_ gene on an IncFIB. Z035 (ST17) and Z017 (ST273) had a *bla*_*VIM*−1_ gene carried on an IncN plasmid, however we did not recover this plasmid in Z017 (ST273) in the long read sequencing data; it may have been lost in culture. Z017 short reads were mapped against the Z035 hybrid assembled and showed that the Z017 IncN plasmid is 0-2 SNPs different from Z035 (ST17), thus a possible plasmid transfer may have occurred between these two isolates. When compared to the database using BLAST, the Z035 IncN plasmid showed only 93% query coverage and 99.6% identity to a single plasmid (CP070567.1) without *bla*_*VIM*−1_ found in a clinical *K. pneumoniae*.

Cluster analysis for plasmids without carbapenemases can be found in Supplementary Fig. 6.

## DISCUSSION

Our study investigated ColR-CRKP isolates from UHC Zagreb. Although we detected common carbapenemases as well as shared plasmid Inc types, when looking in more detail using long read sequencing, we found a complex picture of clonal transmission as well as plasmid exchange (summarized in Fig. 4), involving the dissemination of *bla*_*NDM*−1_, *bla*_*OXA*−48_, and *mcr*-9 resistance genes via clonal and horizontal transfer.

**Figure 4:**
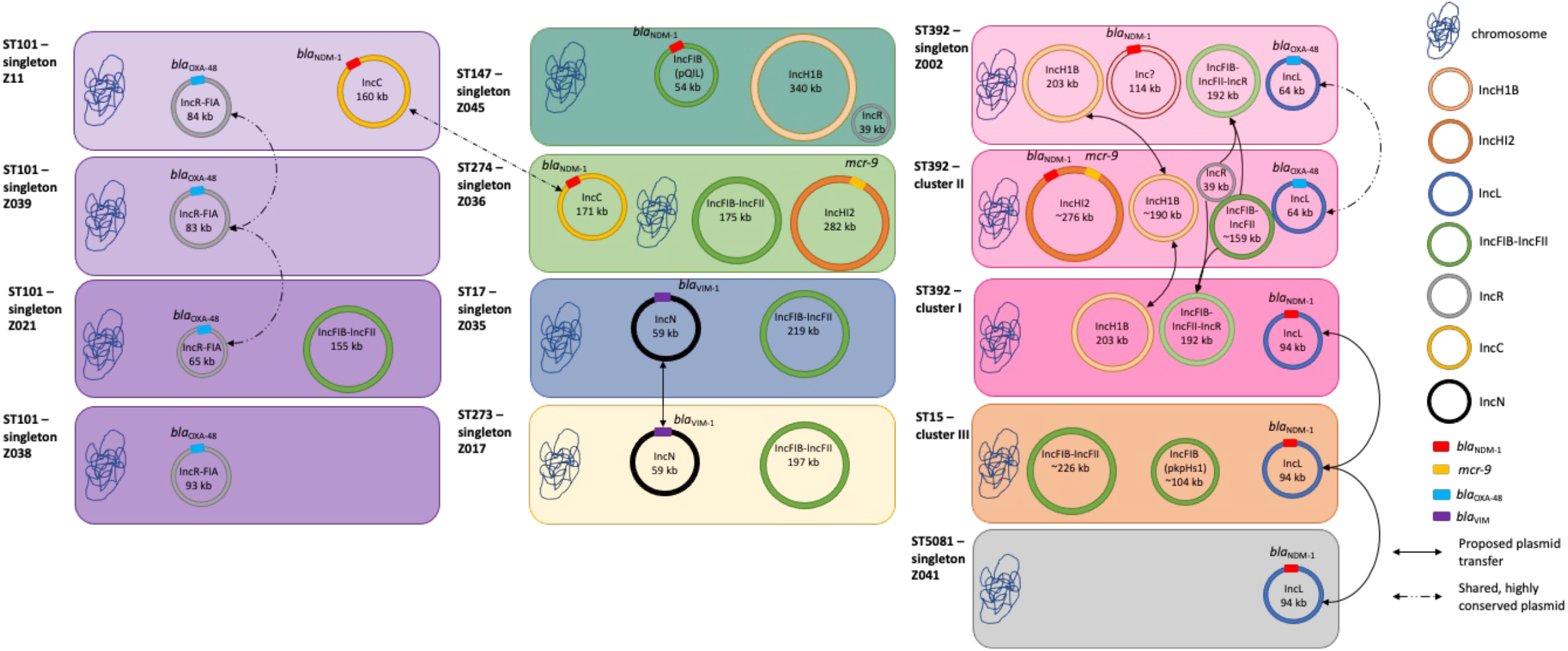
Overview of proposed clonal transmission clusters and horizontal plasmid transfers. Bacterial STs and cluster, plasmid incompatibility groups are highlighted by different colours, as are the relevant antibiotic resistance genes. Proposed horizontal plasmid transfers are indicated by arrows, inconclusive transfers by dashed lines.

From a standard outbreak investigation, we detected three clonal transmissions – two within ST392 (cluster I and cluster II) and one in ST15 (cluster III) (Fig. 1). ST392 has been sporadically identified from hospital-acquired infections, either as a carbapenem resistant and susceptible clone in different parts of Europe, including Spain, Netherlands, United Kingdom, Italy, France, Germany, Luxembourg, Belgium, Türkiye and Austria (36) while ST15 are known to be one of the dominant CRKP lineages of *K. pneumoniae* in clinical samples from European countries, including Croatia, alongside ST11, ST101 and ST258/512 36. These lineages are considered “high risk” clones that have gained a foothold in most parts of southern and Eastern Europe and are most often associated with hospital outbreaks. The earliest report of carbapenemases in Croatia were *bla*_*NDM*_−1 and blaKPC in 2008 and 2012 respectively, detected in *K. pneumoniae* isolates from UHC Zagreb patients (37-38).

From a plasmid perspective, we identified two major IncL plasmids – the promiscuous pOXA-48 (IncL-64kb) shared between ST392 cluster II and ST392 singleton, and a novel 96kb plasmid (IncL-96kb) shared between ST392 cluster I, ST15 cluster III, and ST5081 singleton (Fig 3, Fig. 4). The IncL/M plasmid is known for its worldwide dissemination of *bla*_*OXA*−48_ and its emergence with *bla*_*NDM*−1_ therefore makes it a great public health concern (39). Recently, the IncL/M plasmid group has been re-classified into separate IncL and IncM plasmids, where *bla*_*OXA*−48_ are carried on the IncL plasmid and *bla*_*NDM*−1_ carried on an IncM plasmid (40). In our study, however, we found an IncL pOXA-48 backbone (IncL-64kb) fused with an additional region that contains numerous AMR genes including *bla*_*NDM*−1_ (Supplementary Fig. 2. While mobile genetic elements such as plasmids harbouring antibiotic resistance genes are known to undergo significant gene rearrangement (41), our study shows high stability of this novel IncL-96kb plasmid even when transferred within and between clusters and STs. Given the conserved nature of pOXA-48 and geographically widespread reports of it, we cannot rule out independent acquisition of this plasmid IncL-64kb in our two ST392 clusters. In contrast, given that the novel IncL-96kb has not been described elsewhere and it is highly conserved, this makes horizontal transfer locally between the different STs very likely. Other plasmids carrying *bla*_*OXA*−48_ are less common but have been described, like the IncR-FIA (42) we also found in this collection, as well as an IncFII plasmid (43). The IncN plasmid was also likely shared between two STs, but we were unable to recover it in the long read sequencing. An IncC *bla*_*NDM*−1_ plasmid was observed in two STs, but as we found identical plasmids in the public databases, we cannot ascertain whether it was truly shared or acquired independently. IncC has been reported to play important role in the rapid dissemination of carbapenemases, with *bla*_*NDM*−1_ commonly carried on this plasmid, and has been seen in several bacteria species including Klebsiella species as seen in our study (44).

From the perspective of the carbapenemases, we also find a complex pattern. Apart from the IncL-96kb plasmid described above, we found *bla*_*NDM*−1_ on a 276kb IncHI2, and on a 163kb IncC (Supplementary Table S2, Fig. 4). In IncHI2, the carbapenemase was found together with *mcr*-9 (Supplementary Fig. 3). This plasmid replicon type has been identified as a carrier of multiple resistance genes and also due to its size was termed as a superplasmid. This plasmid is not limited to *K. pneumoniae* (45), and are reported to be key drivers for the spread of *mcr*-9 in Enterobacterales, including *K. pneumoniae* (46). *bla*_*OXA*−48_ was found on the typical IncL-64kb pOXA-48, but also on a several IncR-FIA-type plasmids.

In terms of resistance, all isolates collected as part of this study were highly resistant, we detected a carbapenemase in all of the isolates (Supplementary Table S1). Although all isolates were classed as phenotypically resistant to colistin, only eleven isolates harboured a known genotypic resistance mechanism to colistin (*mcr*-9 and/or pmrB_R256G). This suggests that there are unknown genotypic mechanisms and/or chromosomal mutations contributing to colistin resistance (47). The genetic determinant for colistin resistance, *mcr*-9, was present in five ST392 cluster II isolates as well as one ST274 isolate. Interestingly, a previous study observed the co-localization of *mcr*-9 and carbapenemase genes (*bla*_*NDM*−1_, *bla*_*VIM*−1_ and *bla*_*OXA*−48_) in 26 *K. pneumoniae* isolates belonging to ST274 and ST147 from patients in eight European countries (48), with ST147 being a single locus variant of ST392.

Overall, whilst we observed good overlap of patients in the same wards at the same time, we do not have epidemiological support for all clonal transmissions and none for the proposed horizontal plasmid transfer. There might be intermediate patients that were not screened and where colonization might have gone undetected, and who might be able to close these transmission gaps. Further, only few colonies or single colony picks were selected for each patient, thus colonization with more than one clone of a particular species might have gone undetected. Additionally, for this study, only carbapenem- and colistin-resistant *K. pneumoniae* were considered, missing out on colistin susceptible *K. pneumoniae* as well as other Enterobacterales that might harbour the plasmids under investigation. Lastly, environmental samples were not considered either.

In conclusion, we provide conclusive evidence for clonal transmission of colistin and carbapenem resistant *K. pneumoniae* in Croatia, as well as horizontal plasmid transfer between different ST. Mobile genetic elements have been a driving force in the dissemination of antimicrobial resistance genes most especially in healthcare associated bacterial infections (49), however tracing the transfer of plasmids has been challenging. Short-read sequencing offers limited resolution and would have mislead us in our current study setting: Whilst we detected numerous isolates of the same ST, and both IncL and *bla*_*NDM*−1_ as well as *bla*_*OXA*−48_ and *mcr*-9 in a number of our isolates, the actual picture of spread and dissemination was more complex. Our study therefore emphasizes the importance of bacterial whole genome surveillance with both short- and long-read sequencing as a critical tool for detecting clonal and horizontal transmission of extremely drug-resistant pathogens and then devising appropriate infection prevention and control strategies accordingly.

## ACKNOWLEDGMENTS

We would like to thank Leonardo Duarte dos Santos for excellent technical assistance.

## FIGURE LEGENDS

**Supplementary Table S1: Metadata of collection**. Information on clinical metadata, clustering and sequencing statistics, phenotypic characteristics, identified antibiotic resistance genes and mutations, as well as detected plasmid incompatibility groups.

**Supplementary Table S2: Overview of all detected plasmids, their incompatibility groups, sizes and colistin and/or carbapenem resistance (COL-CR) genes detected**. Same colour in a column indicates plasmid similarities according to mash tree distance.

**Supplementary Figure 1: Spatial and temporal relationship of all patients within the hospital**.

**Supplementary Figure 2: Circular plasmid map of the novel IncL-96kb plasmid**. The tracks from the outside to inside represent: (1) Forward Coding Sequence; (2) Reverse Coding Sequence; (3)

**Supplementary Figure 3: Structural and mutational analysis of IncHI2 plasmids**. (a) Structural relationship among the plasmids analysed using the pling tool, represented as a network. Each plasmid is a node. Edges are labelled with two numbers: the “containment distance” (the proportion of the smaller plasmid which is not alignable to the larger - a low distance means almost all of the small plasmid is contained in the larger) and the “double-cut and join indel distance” (the number of structural rearrangement events, including indels, separating the two genomes). (b) Linear comparison of representative of ST392-cluster II isolates and ST274 (Z036). (c) Detailed view showing the region not found in Z005 IncHI2 plasmid, and (d) showing the region not found in Z003 IncHI2 plasmid. The grey area indicates regions of shared similarities. Red arrows indicate resistance genes, blue arrow indicates transposons, orange arrows indicate the conjugal transfer genes and black arrows are other genes like hypothetical or unknown genes. rep: replication system.

**Supplementary Figure 4: Comparative genomic analysis of IncC plasmid**. (a) Linear comparison of Z011 (ST101) and Z036 (ST274) and (b) detailed view showing the region not found in Z036 isolates which comprises many hypothetical proteins.

**Supplementary Figure 5: Pling relatedness network of IncR-FIA plasmids shows differences in gene content and organisation**. Apart from Z038, other plasmids had 99-100% shared sequence content but are separated by 5-7 rearrangement events.

**Supplementary Figure 6: Structural comparative genomics of the IncHI1B and IncFIB-FII-R plasmids not carrying carbapenemases**. (a) All IncHI1B plasmids (purple) except Z045 (blue) have a containment distance of 0 to 0.01, meaning that they are >99% alignable, with very little gene gain/loss. In addition, the plasmids are very similar in terms of synteny, with 1 to 3 rearrangement events separating them. (b) IncR, IncFIB-FII, and IncFIB-FII-R fusion plasmids. IncR plasmids (highlighted in blue) as well as IncFIB-FII plasmids (red) are very similar within their gropus, with little to no rearrangements. IncFIB-FII-R plasmids (purple) are contained within both groups as indicated by the low containment distances (98-100% shared sequence) and separated by 0-4 rearrangement events. This indicates a fusion of these two separate plasmids, Z002 (ST392 singleton), Z047 and Z013 (both ST392 cluster I). As these isolates come temporally later than the others (all of ST392 cluster II), we assume a fusion event to be most likely as opposed to a split into two plasmids.

## REFERENCES

1. 2015. WHO Library Cataloguing-in-Publication Data Global Action Plan on Antimicrobial Resistance.

2. Ecdc. 2014. SURVEILLANCE REPORT: Antimicrobial resistance surveillance in Europe 2014 10.2900/23549.

3. Pourgholi L, Farhadinia H, Hosseindokht M, Ziaee S, Nosrati R, Nosrati M, Boroumand M. 2022. Analysis of carbapenemases genes of carbapenem-resistant Klebsiella pneumoniae isolated from Tehran heart center.

4. Centers for Disease Control (CDC). 2019. Antibiotic Resistance Threats in the United States, 2019 10.15620/cdc:82532.

5. Al-Farsi HM, Camporeale A, Ininbergs K, Al-Azri S, Al-Muharrmi Z, Al-Jardani A, Giske CG. 2020. Clinical and molecular characteristics of carbapenem non-susceptible Escherichia coli: A nationwide survey from Oman. PLoS One 15:1–23.

6. Nordmann P. 2014. Carbapenemase-producing Enterobacteriaceae: Overview of a major public health challenge. Med Mal Infect 44:51–56.

7. Ecdc. 2019. Carbapenem-resistant Enterobacteriaceae-second update Event background Current situation of CRE in EU/EEA countries.

8. D’Onofrio V, Conzemius R, Varda-Brkić D, Bogdan M, Grisold A, Gyssens IC, Bedenić B, Barišić I. 2020. Epidemiology of colistin-resistant, carbapenemase-producing Enterobacteriaceae and Acinetobacter baumannii in Croatia. Infection, Genetics and Evolution 81.

9. Petrosillo N, Taglietti F, Granata G. 2019. Clinical Medicine Treatment Options for Colistin Resistant Klebsiella pneumoniae: Present and Future 10.3390/jcm8070934.

10. Hammood Hussein N, S AL-Kadmy IM, Mohammed Taha B, Dakel Hussein J. 2021. Mobilized colistin resistance (mcr) genes from 1 to 10: a comprehensive review. Mol Biol Rep 48:2897–2907.

11. Reyes JA, Melano R, Cárdenas PA, Trueba G. 2020. Mobile genetic elements associated with carbapenemase genes in South American Enterobacterales. The Brazilian Journal of Infectious Diseases 24:231–238.

12. Sheppard AE, Stoesser N, Wilson DJ, Sebra R, Kasarskis A, Anson LW, Giess A, Pankhurst LJ, Vaughan A, Grim CJ, Cox HL, Yeh AJ, Sifri D, Walker AS, Peto TE, Crook DW, Mathers AJ. 2016. Nested Russian Doll-Like Genetic Mobility Drives Rapid Dissemination of the Carbapenem Resistance Gene bla KPC 10.1128/AAC.00464-16.

13. De Man TJB, Yaffee AQ, Zhu W, Batra D, Alyanak E, Rowe LA, McAllister G, Moulton-Meissner H, Boyd S, Flinchum A, Slayton RB, Hancock S, Spalding Walters M, Laufer Halpin A, Rasheed JK, Noble-Wang J, Kallen AJ, Limbago BM. 2021. Multispecies Outbreak of Verona Integron-Encoded Metallo-ß-Lactamase-Producing Multidrug-Resistant Bacteria Driven by a Promiscuous Incompatibility Group A/C2 Plasmid. Clinical Infectious Diseases 72:414–420.

14. Bankevich A, Nurk S, Antipov D, Gurevich AA, Dvorkin M, Kulikov AS, Lesin VM, Nikolenko SI, Pham S, Prjibelski AD, Pyshkin A V., Sirotkin A V., Vyahhi N, Tesler G, Alekseyev MA, Pevzner PA. 2012. SPAdes: A New Genome Assembly Algorithm and Its Applications to Single-Cell Sequencing. Journal of Computational Biology 19:455–477.

15. Seemann T. mlst. https://github.com/tseemann/mlst.

16. Ponstingl H, Ning Z. 2010. SMALT-a new mapper for DNA sequencing reads. F1000 Posters, 1(L313).

17. Wood DE, Salzberg SL. 2014. Kraken: Ultrafast metagenomic sequence classification using exact alignments. Genome Biol 15:1–12.

18. Wick RR. 2018. Porechop. San Francisco, CA: GitHub.

19. Wick RR. 2018. Filtlong. San Francisco, CA: GitHub.

20. Wick RR, Judd LM, Gorrie CL, Holt KE. 2017. Unicycler: Resolving bacterial genome assemblies from short and long sequencing reads. PLoS Comput Biol 13:e1005595.

21. Mikheenko A, Prjibelski A, Saveliev V, Antipov D, Gurevich A. 2018. Versatile genome assembly evaluation with QUAST-LG. Bioinformatics 34:i142–i150.

22. Auwera GAV der. 2020. Genomics in the Cloud: Using Docker, GATK, and WDL in Terra. O’Reilly, Sebastopol.

23. Croucher NJ, Page AJ, Connor TR, Delaney AJ, Keane JA, Bentley SD, Parkhill J, Harris SR. 2015. Rapid phylogenetic analysis of large samples of recombinant bacterial whole genome sequences using Gubbins. Nucleic Acids Res 43:15.

24. Letunic I, Bork P. 2021. Interactive Tree Of Life (iTOL) v5: an online tool for phylogenetic tree display and annotation. Nucleic Acids Res 49:W293–W296.

25. Zankari E, Hasman H, Cosentino S, Vestergaard M, Rasmussen S, Lund O, Aarestrup FM, Larsen M V. 2012. Identification of acquired antimicrobial resistance genes. Journal of Antimicrobial Chemotherapy 67:2640–2644.

26. Carattoli A. 2009. Resistance plasmid families in Enterobacteriaceae. Antimicrob Agents Chemother 10.1128/AAC.01707-08.

27. Schwengers O, Jelonek L, Dieckmann MA, Beyvers S, Blom J, Goesmann A. 2021. Bakta: Rapid and standardized annotation of bacterial genomes via alignment-free sequence identification. Microb Genom 7:000685.

28. Ondov BD, Treangen TJ, Melsted P, Mallonee AB, Bergman NH, Koren S, Phillippy AM. 2016. Mash: Fast genome and metagenome distance estimation using MinHash. Genome Biol 17.

29. Roberts LW, Enoch DA, Khokhar F, Blackwell GA, Wilson H, Warne B, Gouliouris T, Iqbal Z, Török ME. 2023. Long-read sequencing reveals genomic diversity and associated plasmid movement of carbapenemase-producing bacteria in a UK hospital over 6 years. Microb Genom 9.

30. Madden T. 2002. Chapter 16: The BLAST Sequence Analysis Tool. The NCBI Handbook[internet].

31. Sullivan MJ, Petty NK, Beatson SA. 2011. Easyfig: A genome comparison visualizer. Bioinformatics 27.

32. Carver T, Thomson N, Bleasby A, Berriman M, Parkhill J. 2009. DNAPlotter: circular and linear interactive genome visualization. Bioinformatics 25:119–120.

33. Schmartz GP, Hartung A, Hirsch P, Kern F, Fehlmann T, Müller R, Keller A. 2022. PLSDB: advancing a comprehensive database of bacterial plasmids. Nucleic Acids Res 50:D273–D278.

34. Hawkey J, Wyres KL, Judd LM, Harshegyi T, Blakeway L, Wick RR, Jenney AWJ, Holt KE. 2022. ESBL plasmids in Klebsiella pneumoniae: diversity, transmission and contribution to infection burden in the hospital setting. Genome Med 14:97.

35. Shannon P, Markiel A, Ozier O, Baliga NS, Wang JT, Ramage D, Amin N, Schwikowski B, Ideker T. 2003. Cytoscape: A Software Environment for Integrated Models of Biomolecular Interaction Networks. Genome Res 13:2498–2504.

36. David S, Reuter S, Harris SR, Glasner C, Feltwell T, Argimon S, Abudahab K, Goater R, Giani T, Errico G, Aspbury M, Sjunnebo S, Koraqi A, Lacej D, Apfalter P, Hartl R, Glupczynski Y, Huang TD, Strateva T, Marteva-Proevska Y, Andrasevic AT, Butic I, Pieridou-Bagatzouni D, Maikanti-Charalampous P, Hrabak J, Zemlickova H, Hammerum A, Jakobsen L, Ivanova M, Pavelkovich A, Jalava J, Österblad M, Dortet L, Vaux S, Kaase M, Gatermann SG, Vatopoulos A, Tryfinopoulou K, Tóth Á, Jánvári L, Boo TW, McGrath E, Carmeli Y, Adler A, Pantosti A, Monaco M, Raka L, Kurti A, Balode A, Saule M, Miciuleviciene J, Mierauskaite A, Perrin-Weniger M, Reichert P, Nestorova N, Debattista S, Mijovic G, Lopicic M, Samuelsen Ø, Haldorsen B, Zabicka D, Literacka E, Caniça M, Manageiro V, Kaftandzieva A, Trajkovska-Dokic E, Damian M, Lixandru B, Jelesic Z, Trudic A, Niks M, Schreterova E, Pirs M, Cerar T, Oteo J, Aracil B, Giske C, Sjöström K, Gür D, Cakar A, Woodford N, Hopkins K, Wiuff C, Brown DJ, Feil EJ, Rossolini GM, Aanensen DM, Grundmann H. 2019. Epidemic of carbapenem-resistant Klebsiella pneumoniae in Europe is driven by nosocomial spread. Nat Microbiol 4:1919–1929.

37. Bedenić B, Mazzariol A, Plečko V, Bošnjak Z, Barl P, Vraneš J, Cornaglia G. 2012. First report of KPC-producing Klebsiella pneumoniae in Croatia. Journal of Chemotherapy 24:237–239.

38. Mazzariol A, Bošnjak Z, Ballarini P, Budimir A, Bedenić B, Kalenić S, Cornaglia G. 2012. NDM-1–producing Klebsiella pneumoniae, Croatia. Emerg Infect Dis 18:532.

39. Getino M, López-Díaz M, Ellaby N, Clark J, Ellington MJ, La Ragione RM. 2022. A Broad-Host-Range Plasmid Outbreak: Dynamics of IncL/M Plasmids Transferring Carbapenemase Genes. Antibiotics 11:1641.

40. Carattoli A, Seiffert SN, Schwendener S, Perreten V, Endimiani A. 2015. Differentiation of IncL and IncM Plasmids Associated with the Spread of Clinically Relevant Antimicrobial Resistance. PLoS One 10:e0123063.

41. Mathers AJ, Crook D, Vaughan A, Barry KE, Vegesana K, Stoesser N, Parikh HI, Sebra R, Kotay S, Walker AS, Sheppard AE. 2019. Klebsiella quasipneumoniae Provides a Window into Carbapenemase Gene Transfer, Plasmid Rearrangements, and Patient Interactions with the Hospital Environment. Antimicrob Agents Chemother 63.

42. Campos-Madueno EI, Moser AI, Jost G, Maffioli C, Bodmer T, Perreten V, Endimiani A. 2022. Carbapenemase-producing Klebsiella pneumoniae strains in Switzerland: human and non-human settings may share high-risk clones. J Glob Antimicrob Resist 28:206–215.

43. Moussa J, Panossian B, Nassour E, Salloum T, Abboud E, Tokajian S. 2020. Detailed characterization of an IncFII plasmid carrying blaOXA-48 from Lebanon. Journal of Antimicrobial Chemotherapy 75:2462–2465.

44. Carattoli A, Villa L, Poirel L, Bonnin RA, Nordmann P. 2012. Evolution of IncA/C bla CMY-2-Carrying Plasmids by Acquisition of the bla NDM-1 Carbapenemase Gene. Antimicrob Agents Chemother 56:783–786.

45. García-Fernández A, Carattoli A. 2010. Plasmid double locus sequence typing for IncHI2 plasmids, a subtyping scheme for the characterization of IncHI2 plasmids carrying extended-spectrum β-lactamase and quinolone resistance genes. Journal of Antimicrobial Chemotherapy 65:1155–1161.

46. Macesic N, Blakeway L V, Stewart JD, Hawkey J, Wyres KL, Judd LM, Wick RR, Jenney AW, Holt KE, Peleg AY. 2021. Silent spread of mobile colistin resistance gene mcr-9.1 on IncHI2 “superplasmids” in clinical carbapenem-resistant Enterobacterales. Clin Microbiol Infect 27.

47. Hinchliffe P, Yang QE, Portal E, Young T, Li H, Tooke CL, Carvalho MJ, Paterson NG, Brem J, Niumsup PR, Tansawai U, Lei L, Li M, Shen Z, Wang Y, Schofield CJ, Mulholland AJ, Shen J, Fey N, Walsh TR, Spencer J. 2017. Insights into the Mechanistic Basis of Plasmid-Mediated Colistin Resistance from Crystal Structures of the Catalytic Domain of MCR-1. Sci Rep 7:39392.

48. Wang Y, Liu F, Hu Y, Zhang G, Zhu B, Gao GF. 2020. Detection of mobile colistin resistance gene mcr-9 in carbapenem-resistant Klebsiella pneumoniae strains of human origin in Europe. Journal of Infection 80:578–606.

49. Lerminiaux NA, Cameron ADS. 2019. Horizontal transfer of antibiotic resistance genes in clinical environments. Can J Microbiol 65:34–44.

50. Frolova D, Lima L, Roberts L, Bohnenkämper L, Wittler R, Stoye J, Iqbal Z. 2024. Applying rearrangement distances to enable plasmid epidemiology with pling. bioRxiv 10.1101/2024.06.12.598623v1.

